# DNMT family induced down-regulation of NDRG1 via DNA methylation and clinicopathological significance in gastric cancer

**DOI:** 10.1101/2021.03.01.433329

**Authors:** Xiaojing Chang, Jinguo MA, Xiaoying Xue, Guohui Wang, Linlin Su, Xuetao Han, Huandi Zhou, Liubing Hou

## Abstract

**Background:** Aberrant DNA methylation of tumor suppressor genes is a common event in the development and progression of gastric cancer(GC). Our previous study showed NDRG1, which could suppress cell invasion and migration, was frequently down-regulated by DNA methylation of its promoter in GC.

**Purpose and Methods:** To analyze the relationship between the expression and DNA methylation of NDRG1 and DNA methyltransferase (DNMT) family. We performed a comprehensive comparison analysis using 407 patients including sequencing analysis data of GC from TCGA.

**Results:** NDRG1 was negatively correlative to DNMT1 (*p* =0.03), DNMT3A(*p* =0.01), DNMT3B(*p* =0.88), respectively. Whereas, the DNA methylation of NDRG1 was positively correlative to DNMT family(DNMT1 p<0.01, DNMT3A p<0.001, DNMT3B p=0.57, respectively). NDRG1 expression was significantly inverse correlated with invasion depth (*p* =0.023), and DNMT1 was significantly positive correlated with the degree of tumor cell differentiation (*p* =0.049). DNMT3B was significantly correlated with tumor cell differentiation (*p* =0.030). However, there was no association between the expression of DNMT3A and clinicopathological features. The univariate analysis showed that NDRG1and DNMTs had no association with prognosis of GC patients. But, multivariate analysis showed DNMT1 was significantly correlated with prognosis of GC patients.

**Conclusion:** These data suggest that down-regulation of NDRG1 in gastric cancer is due to DNA methylation of NDRG1 gene promoter via DNMT family. The demethylating agent maybe a potential target drug for GC patients.

## Introduction

Gastric carcinoma (GC) is one of the most common digestive malignancies worldwide with the third most common cause of cancer-related death^1, 2^. The poor prognosis and high recurrence rate are mainly due to lymph node metastasis in its early stage.

To date, a number of laboratories have shown that the occurrence of GC is usually caused by oncogenes activation and tumor suppressor genes(TSGs) inactivation. TSGs could suppress tumor cells migration and invasion. Studies have shown that the inactivation of most TSGs are related to DNA methylation of CpG, while the silencing of these genes possibly contributes to the development and progression of tumors^3-5^. There is increasing evidence that aberrant DNA promoter methylation of TSGs is a major event in the development and progression of gastric cance^6^. Our previous studies showed NDRG1(N-myc downstream-regulated gene 1), a differentiation-related and tumor suppressor gene, which could suppress cancer cells invasion and migration, was frequently down-regulated by DNA methylation of its promoter in GC, but there were not any mutations in NDRG1 cDNA, and also not significant correlation with histone acetylation ^7, 8^.

DNA methylation of CpG islands, which is carried out by DNA methyltransferase (DNMT) enzymes, is the most widely and the best well studied epigenetic modification event, and leads to transcriptional gene silencing^9^. DNA methyltransferase (DNMT) enzymes include DNMT1, DNMT2, DNMT3A, DNMT3B, and DNMT3L, among which DNMT1(a maintenance DNMT), DNMT3A and DNMT3B(de novo methyltransferases) are the most important^10^. In the current study, we focused on the relationship between NDRG1 and DNMT family(DNMT1, DNMT3A and DNMT3B) and their clinical significance in GC by analyzing high-throughput data obtained from TCGA.

## Materials and methods

The data that we investigated the mRNA expression of NDRG1 and DNMTs, and the DNA methylation of NDRG1 in GC was obtained from the data portal of TCGA (https://portal.gdc.cancer.gov/), which is a well-known cancer research project that collects and analyzes high-quality tumor samples and makes the related data available to researchers. We obtained the GC data set (March 1st, 2018 updated), which encompasses 407 gastric tumor samples, and of which 338 cases encompasses the data of DNA methylation of NDRG1. Total of 315 cases contain the complete clinicopathological information.

### Gastric Cells and Culture

Human gastric cancer cell lines, SGC7901 and MKN45 were obtained from the Institute of Biochemistry and Cell Biology, Chinese Academy of Sciences (Shanghai, China), and one immortalized normal gastric cell line, GES1 was obtained from the Oncology Institute of China Medical University. Cells were cultured with RPMI 1640 (Invitrogen/Gibco, Grand Island, NY, USA) containing 10 % fetal bovine serum (Invitrogen/Gibco) in a humidified atmosphere of 5 % CO2 at 37 C.

### Protein Isolation and Western Blot

Total cellular protein was extracted from cultured cells and western bolt was performed as previous description^7^. Protein bands were scanned and quantified using densitometric software (Bio-Rad, California, USA). A rabbit monoclonal anti-NDRG1 antibody was at a dilution of 1:1,000 (Cell Signaling Technology, Danvers, MA, USA). DNMT1, DNMT3A and DNMT3B antibody at a dilution of 1:500 were purchased from Bioss, Beijing, China.

### Statistics

All statistical analyses were conducted using SPSS 24.0 (Chicago, IL, USA) and R (https://www.r-project.org/). R language and GraphPad Prism 7 (San Diego, CA, USA) were performed to draw plots. X^2^ test and Fisher’s exact test were used to generate P values, Values with *p* < 0.05 are considered as statistically significant.

## Results

NDRG1 was negatively correlative to DNMT family

Total of 407 gastric cancer cases (375 was gastric cancer tissues and 32 was nonmalignant gastric tissues) which included sequencing analysis data were downloaded form TCGA, then we performed a comprehensive comparison analysis about the mRNA expression of NDRG1 and DNMTs and their clinicopathological features. As shown in Figure 1A-C, NDRG1 mRNA expression was negatively correlative to DNMT1(r=-0.1067, *p* =0.03), DNMT3A(r=-0.1015, *p* =0.01), DNMT3B(r=-0.0069, *p* =0.88). Noteworthy, NDRG1 was significantly associated with DNMT1(*p* =0.03) and DNMT3A (*p* =0.01), and DNMT3A showed the strongest association. However, there was no significant association between NDRG1 and DNMT3B (*p*=0.8885).

**Figure.1.**
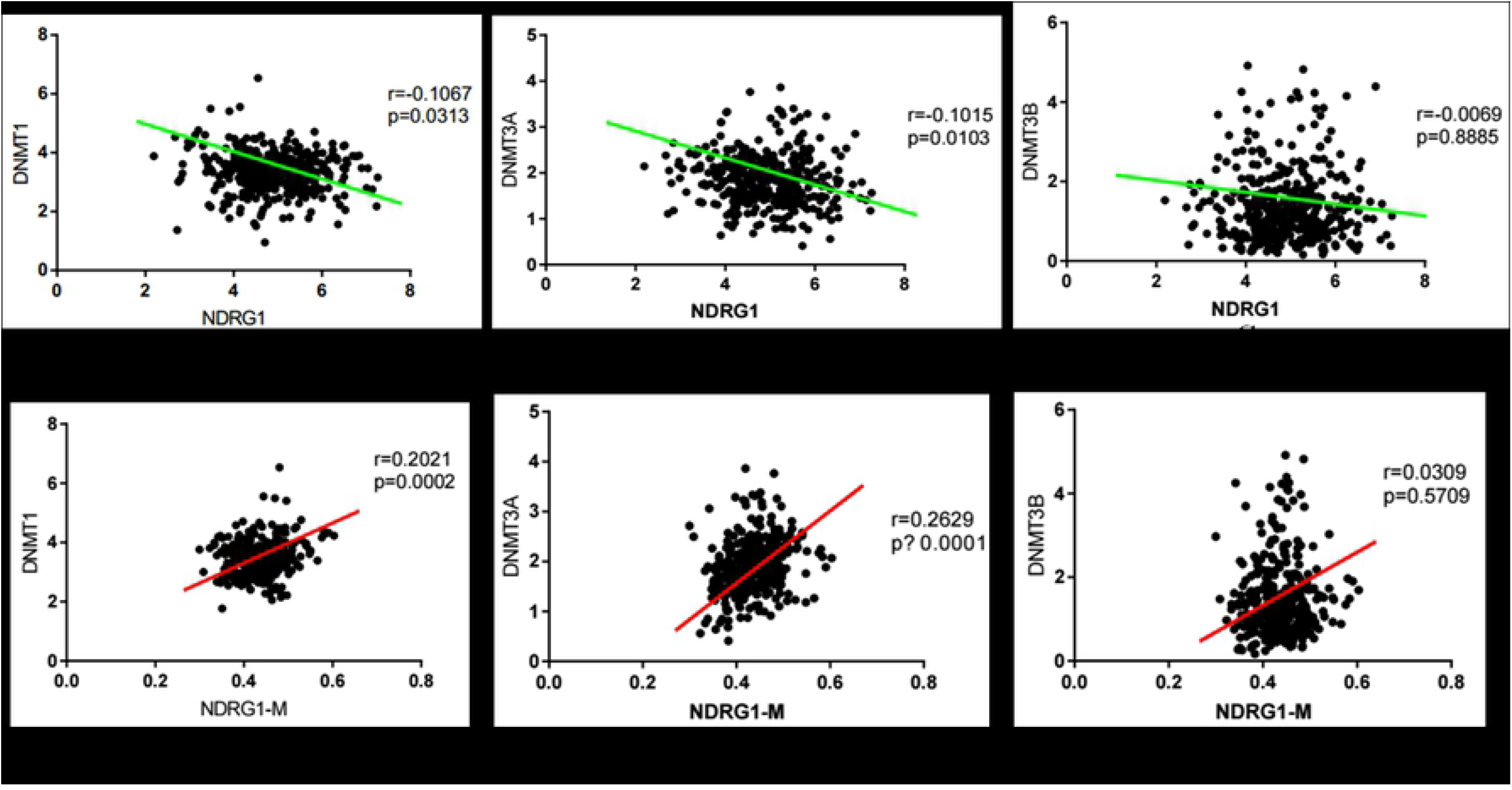
Association between NDRG1 and DNMTs. A-C, NDRG1 mRNA expression was negatively correlative to DNMT1, DNMT3A and DNMT3B. D-F, DNA methylation of NDRG1 gene promoter was positively correlative to the expression DNMT family.

To confirm these results, we further assessed the protein levels of NDRG1 and DNMTs in one immortalized normal gastric cell line, GES1 and two gastric cell lines, SGC7901 and MKN45 by western blot. Results showed DNMT1, DNMT3A and DNMT3B were up-regulated in SGC7901 and MKN45 cells compared with GES1, while NDRG1 expression was inversed (*p* < 0.05, Fig. 2). This was in accordance to the results of the relationship between NDRG1 and DNMT family mRNA expression from TCGA.

**Figure.2.**
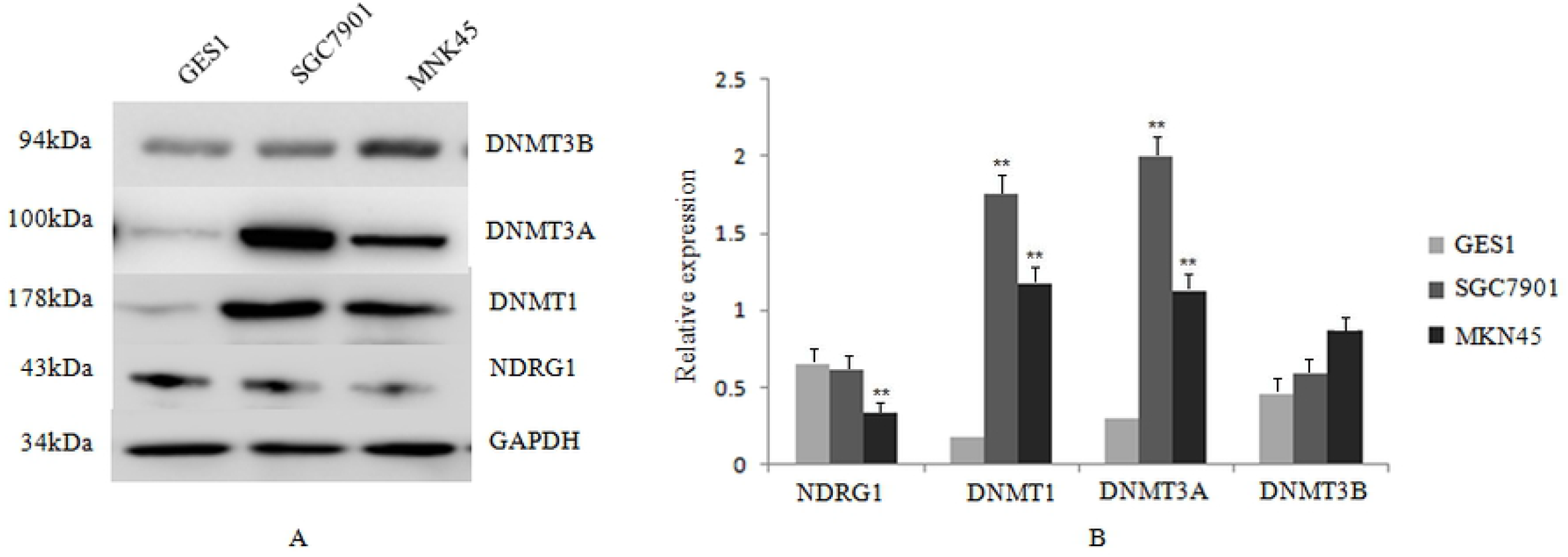
The protein levels of NDRG1 and DNMTs in GES1 and two gastric cell lines, SGC7901 and MKN45.A, the protein level of NDRG1 and DNMTs; B. Histogram of the protein level.

### Methylation of NDRG1 gene promoter was positively correlative to the expression DNMT family

Among all of 407 GC cases, 338 cases encompass the data of DNA methylation of NDRG1. Results showed DNA methylation of NDRG1 gene promoter was positively correlative to the expression DNMT family, and was significantly associated with DNMT1 (*p* =0.002, Fig.1D) and DNMT3A (*p* <0.001, Fig.1E). However, no statistical significance was found between NDRG1 and DNMT3B (*p* =0.57, Fig. 1F).

### The association of NDRG1 and DNMT family with the clinicopathological parameters and prognosis of GC

The association of NDRG1 and DNMT family with the clinicopathological parameters of gastric cancer patients were shown in Table 1. Total of 315 cases which contain the complete clinicopathological and survival information were analyzed. The high and low expression of NDRG1 and DNMTs was based on the median value of their mRNA levels. We found NDRG1 expression was significantly inverse correlated with invasion depth (*p* =0.023), whereas DNMT1 was statistically significantly positive correlated with invasion depth (*p* = 0.049), DNMT3B was significantly positive correlated with the degree of tumor cell differentiation (*p* =0.030). There was no significant association between DNMT3A and the clinicopathological parameters of the GC patients (Table 1).

**Table.1.** Clinicopathological parameters of NDRG1 and DNMTs mRNA expression from TCGA cohort

|  |  |  |  |  |  |  |  |  |  |
| --- | --- | --- | --- | --- | --- | --- | --- | --- | --- |
| Variable | Patients | NDRG1 |  | DNMT1 |  | DNMT3A |  | DNMT3B |  |
|  | (n) | Low(High) | p | Low(High) | p | Low(High) | p | Low(High) | p |
| Gender |  |  |  |  |  |  |  |  |  |
| Female | 120 | 61(59) | 0.908 | 56(64) | 0.355 | 56(64) | 0.355 | 61(59) | 0.817 |
| Male | 195 | 97(98) |  | 102 (93) |  | 102(93) |  | 96(99) |  |
| Age |  |  |  |  |  |  |  |  |  |
| <60 | 100 | 52(48) | 0.717 | 47(53) | 0.469 | 50(50) | 1.000 | 49(51) | 0.904 |
| ≥60 | 215 | 106(109) |  | 111(104) |  | 108(107) |  | 108(107) |  |
| Stage |  |  |  |  |  |  |  |  |  |
| I | 42 | 18(24) | 0.297 | 23(19) | 0.881 | 26(16) | 0.190 | 20(22) | 0.814 |
| II | 101 | 47(54) |  | 51(50) |  | 51(50) |  | 49(52) |  |
| III | 139 | 78(61) |  | 69(70) |  | 62(77) |  | 69(70) |  |
| IV | 33 | 15(18) |  | 15(18) |  | 19(14) |  | 19(14) |  |
| Invasion depth |  |  |  |  |  |  |  |  |  |
| T1 | 15 | 9(6) | 0.023* | 12(3) | 0.049* | 11(4) | 0.090 | 9(6) | 0.216 |
| T2 | 63 | 21(42) |  | 33(30) |  | 37(26) |  | 28(35) |  |
| T3 | 151 | 84(67) |  | 77(74) |  | 69(82) |  | 70(81) |  |
| T4 | 66 | 44(42) |  | 36(50) |  | 41(45) |  | 50(36) |  |
| Lymph node metastasis(N) |  |  |  |  |  |  |  |  |  |
| N0 | 99 | 46(53) | 0.348 | 49(50) | 0.828 | 52(47) | 0.864 | 50(49) | 0.206 |
| N1 | 83 | 48(35) |  | 40(43) |  | 43(40) |  | 44(39) |  |
| N2 | 69 | 31(38) |  | 38(31) |  | 33(36) |  | 27(42) |  |
| N3 | 64 | 33(31) |  | 31(33) |  | 30(34) |  | 36(28) |  |
| Distant metastasis |  |  |  |  |  |  |  |  |  |
| No | 295 | 149(146) | 0.652 | 149(146) | 0.652 | 145(150) | 0.248 | 144(151) | 0.174 |
| Yes | 20 | 9(11) |  | 9(11) |  | 13(7) |  | 13(7) |  |
| Differentiation |  |  |  |  |  |  |  |  |  |
| G1 | 7 | 3(4) | 0.293 | 4(3) | 0.906 | 3(4) | 0.758 | 3(4) | 0.030* |
| G2 | 108 | 48(60) |  | 55(53) |  | 57(51) |  | 43(65) |  |
| G3 | 200 | 107(93) |  | 99(101) |  | 98(102) |  | 111(89) |  |
| *p<0.05 |  |  |  |  |  |  |  |  |  |

Univariate analysis of prognosis showed that age, invasion depth, lymph metastasis and stage were significantly inverse correlated with prognosis of GC patients. But NDRG1and DNMTs showed no association with prognosis of GC patients(Fig. 3). Whereas, Multivariate analysis of prognosis showed age, stage and DNMT1 were significantly correlated with prognosis of GC patients. But NDRG1, DNMT3A andDNMT3B showed no association with prognosis(Fig. 4).

**Figure.3.**
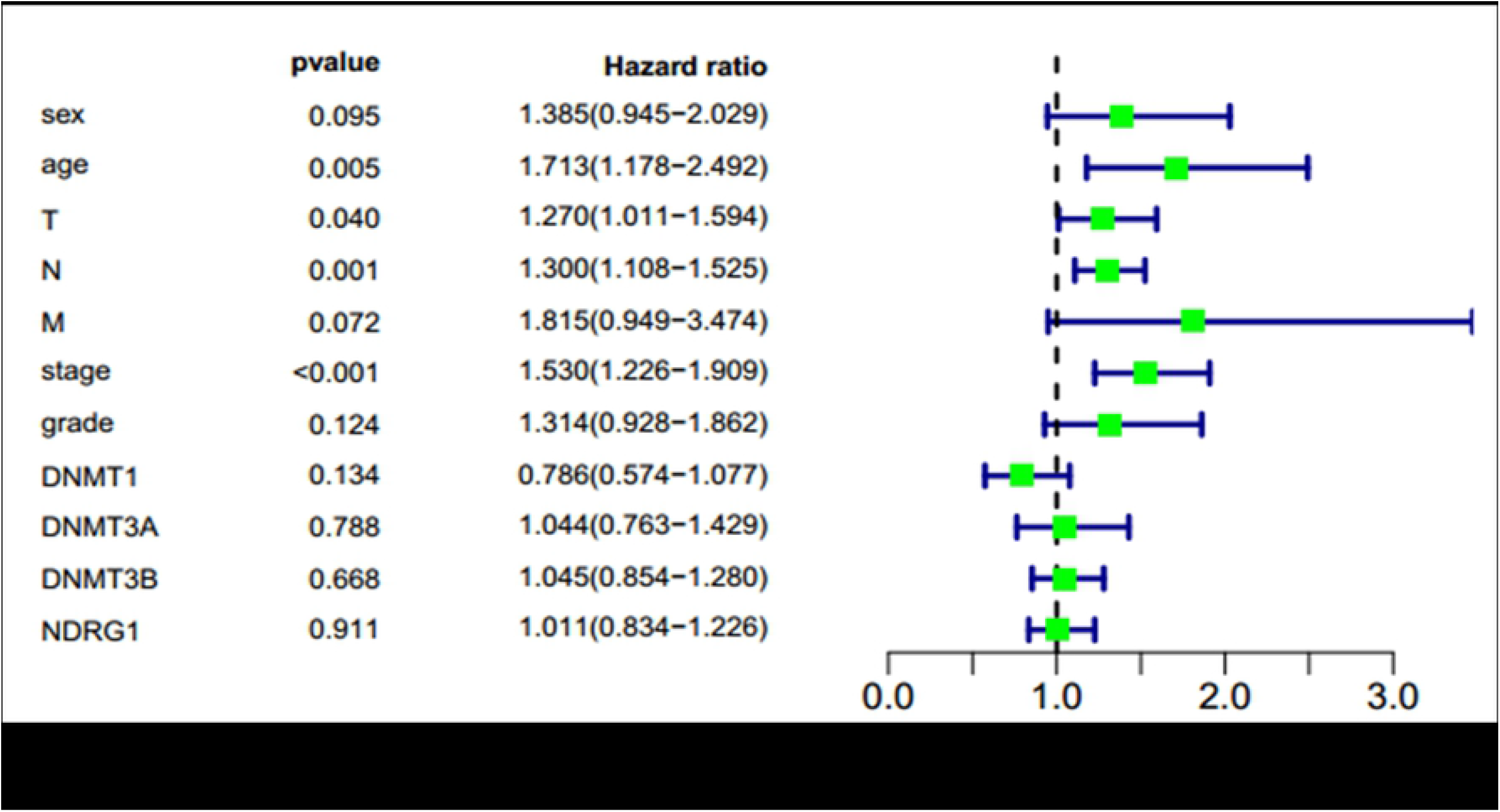
Univariate analysis of prognosis of GC patients.

**Figure.4.**
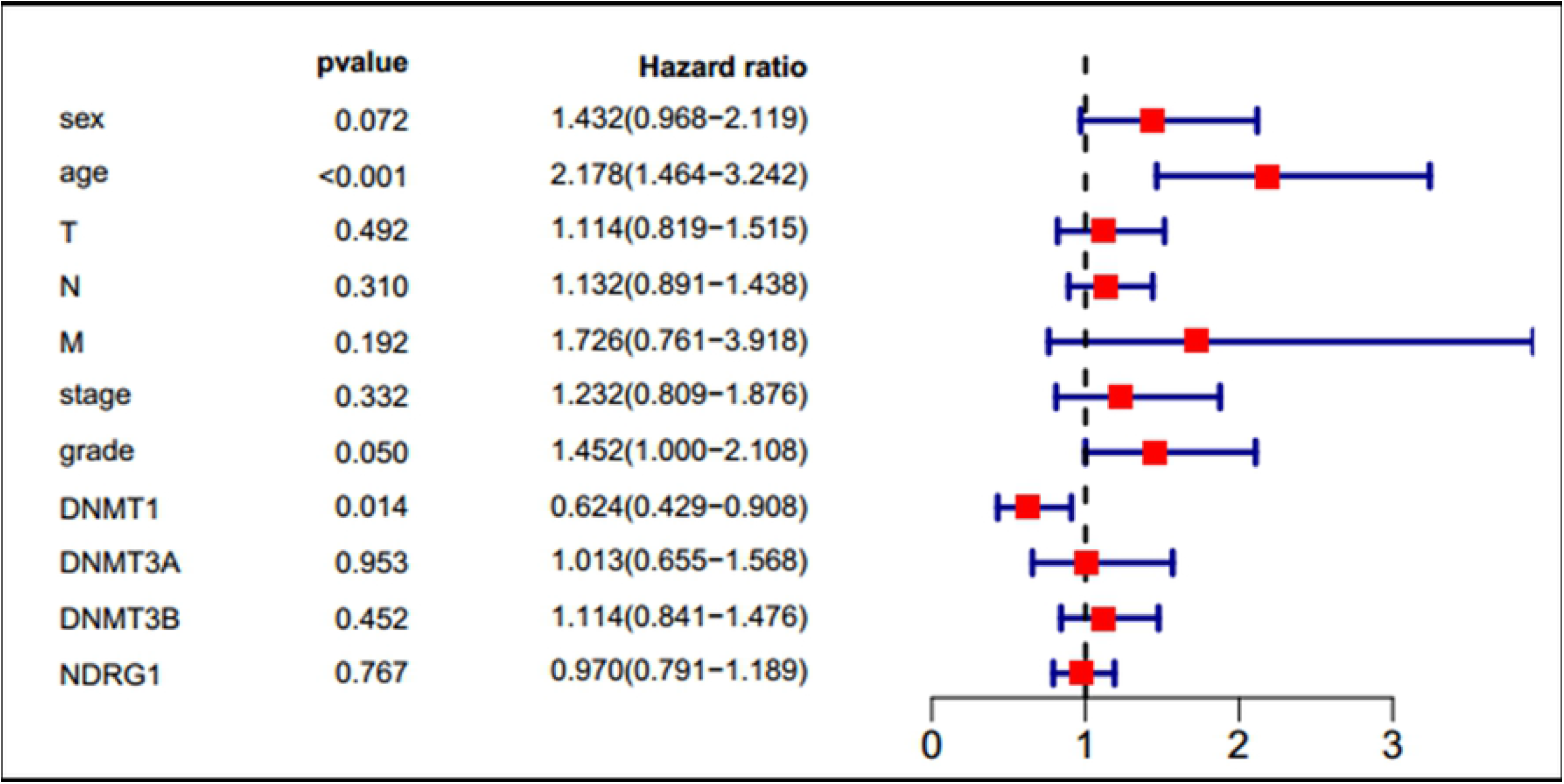
Multivariate analysis of prognosis of GC patients.

## Discussion

NDRG1, a differentiation-related gene, which belongs to NDRG family, is down-regulated in many tumors, and proposed as a metastasis suppressor gene^11-14^. Our previous studies confirmed the anti-cancer effect of NDRG1 in GC^7, 8, 15^. Gene network analysis shows that the 5 ‘end of the NDRG1 gene contains a large CpG island. Some studies have confirmed the downregulation of NDRG1 caused by DNA methylation of CpG islands of its promoter in breast cancer, prostate cancer cells and pancreatic cancer^16-18^. In GC, the down-regulation of NDRG1 was also found to be regulated by DNA methylation of its promoter in our previous study. DNA methylation is carried out by DNA methyltransferase (DNMT) enzymes. To date, not any studies reported the relationship of NDRG1 and DNMTs in GC.

In the current study, total 407 GC cases which included sequencing analysis data of NDRG1 mRNA were analyzed. We found that NDRG1 mRNA level was negatively correlative to DNMT1, DNMT3A and DNMT3B, and DNMT3A showed the strongest association. No significant association was found between NDRG1 and DNMT3B. The results of western blot analysis confirmed these data from TCGA. This is the first time for analyzing the relationship between NDRG1 and DNMT family. NDRG2, another member of NDRG family which includes four members NDRG1-4, was reported to be significantly decreased in human cancers, and in GC, H. pylori silenced Ndrg2 by activating the NF-κB pathway and up-regulating DNMT3b, promoting gastric cancer progression^19^. Our current data were similar to those of NDRG2. Noteworthy, our data showed that aberrant DNA methylation of NDRG1was mainly due to DNMT3A.

We further investigate the association of DNA methylation of NDRG1 and DNMTs. Total 338 of all 407 GC cases encompasses the data of DNA methylation of NDRG1. Conversely, DNA methylation of NDRG1 was positively correlative to the expression of DNMTs, and was significantly associated with DNMT1 and DNMT3A.These results once again confirmed the relationship of NDRG1 mRNA expression and DNMTs in GC. Whereas, NDRG1 expression was significantly inverse correlated with invasion depth, and DNMT1 was positively correlated with invasion depth. DNMT3B was significantly positive correlated with the degree of tumor cell differentiation. But, the univariate analysis of prognosis showed no association between NDRG1and DNMTs with the prognosis of GC patients. While, multivariate analysis of prognosis showed only DNMT1 was significantly correlated with prognosis of GC patients. These results were mainly based on the data of bioinformatics, the current data remain suggest that high NDRG1 expression and low DNMTs expression could suppress cell invasion and migration, may improved the rate of recurrence and metastasis. Our previous study had showed GC patients with high expression of NDRG1 had better overall survival rate than those with low NDRG1 expression^8^. So large sample data of immunohistochemistry, western blot and RT-PCR of GC tissues were urgently needed. DNMT inhibitor, including azacitidine, 5-aza-2′-deoxycytidine (5-Aza-Dc, decitabine), guadecitabine, hydralazine, procaine, MG98 and/or zebularine, among which 5-Aza-Dc is the most common used, could cause demethylating and reactivate the expression of TSGs, then suppress the metastasis of tumor cell^20-22^. NDRG1 expression was found to be increased after treatment with 5-Aza-Dc in breast cancer, prostate cancer and pancreatic cancer^16-18^. Lee E et al^23^ found that the epidrug 5-Aza could up-regulated NDRG1 expression by reduction of suppressive histone marks, H3K9me3 and H3K27me3 on NDRG1 promoter in prostate cancer.

In the current study, the aberrant DNA methylation of NDRG1was found to be significantly associated with DNMT family, especially DNMT1 and DNMT3A. Our previous study showed that 5-Aza-Dc could up-regulate the expression of NDRG1 in GC cells. It suggests that the demethylating agent maybe a potential target drug in GC. Up to now, demethylating treatment has been used for the treatment of myeloid leukemias^24, 25^. In vitro studies of solid tumors, demethylating agents combined chemotherapy and/or immunotherapy could enhances the therapeutic effect in cancer cells^26-29^. In a case report, azacitidine was used to treat a 57-year-old woman newly diagnosed with MDS during palliative chemotherapy for metastatic breast cancer, azacitidine showed promising effects for MDS, and also stabilized the patient’s lung and lymph node metastases without any major toxicity^30^. Now, An increasing number of clinical trials of demethylating treatment is ongoing, some have presented exciting effects^31-34^(Table 2). A Phase I study showed the security of decitabine treated by hepatic arterial infusionin, and a dose level of 20 mg/m2/day on five consecutive days every 4 weeks could be considered for further investigation in combinatorial immunotherapy regimens in patients with Unresectable Liver-predominant Metastases From Colorectal Cancer^33^. In a phase I/II clinical trial, 15 patients with metastatic castration-resistant prostate cancer (mCRPC) were enrolled in phase I which is dose climbing experiment of 5-Aza-Dc, and 7 patients were enrolled in phase II. In phase I, no dose-limiting toxicity (DLT) was observed, and most patients were well tolerated. The highest level reached was 5-Aza-Dc with 150 mg/m2 daily for 5 days followed by docetaxel with 75 mg/m2 on day 6, which was considered as the recommended phase II dose. In phase II, 6 patients received the recommended phase II dose during 46 cycles. Two episodes of Grade 3 hematologic and three Grade 3 nonhematologic toxicities were observed, and one patient died from neutropenic sepsis. Subsequently, 5-Aza was reduced to 75 mg/m2, and no treatment-related>Grade 3 toxicities was observed in one patient. In this clinical trial, PSA response was observed in 10 of 19 (52.6 %) patients, and the median duration of response was 20.5 weeks. Kaplane-Meier estimate of median PFS was 4.9 months, and median OS was 19.5 months, which were both favorable^34^. It suggests that the combination of azacitidine and chemotherapy is active in mCRPC patients, and it maybe a new treatment in future for tumors.

**Table.2.**
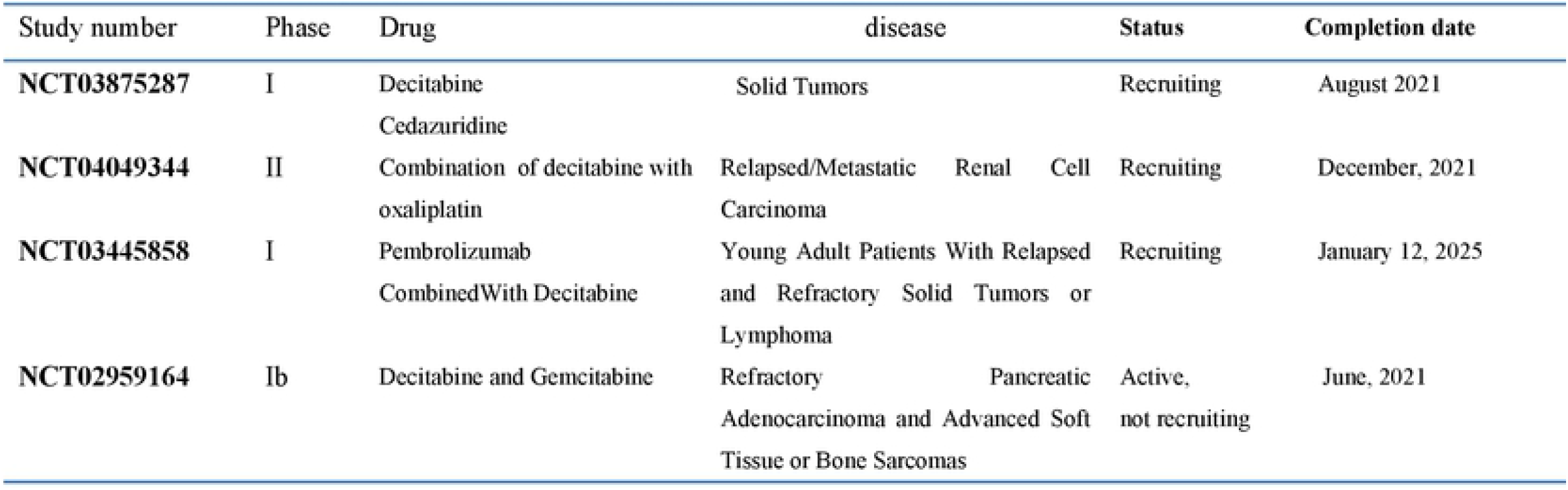
Ongoing clinical trials evaluating decitabine drug in solid tumors

This study is mainly based on the data of bioinformatics, large sample data of immunohistochemistry, western blot and RT-PCR of GC tissues were urgently needed.

We will further detect and confirm the relationship of NDRG1 and DNMTs in GC tissues.

In conclusion, the current data showed that aberrant DNA methylation of NDRG1 which could suppress cell invasion and migration, was regulated by DNMT family in GC, especially DNMT1 and DNMT3A. Furthermore, the demethylating agent 5-Aza-Dc, a DNMT inhibitor, maybe a potential target drug. Therefore, further clinical studies are warranted to evaluate and confirm the effect of demethylating treatment in GC.

## Acknowledgments

This study was supported in part by a grant from the National Natural Science Foundation of Hebei province of China(#H2018206180).

